# Biological motion as an innate perceptual mechanism driving social affiliation

**DOI:** 10.1101/347419

**Authors:** Johannes Larsch, Herwig Baier

## Abstract

Collective behavior, such as shoaling in teleost fish, is driven by the perceptual recognition of conspecific animals. Because social interactions are mutual, it has been difficult to disentangle the exact sensory cues that trigger affiliation in the first place from those that are emitted by receptive and responsive shoal mates. Here we overcome this challenge in a virtual reality assay in zebrafish. We discovered that simple visual features of conspecific biological motion provide an irresistible shoaling cue. Individual juvenile fish interact with circular black dots projected onto a screen, to the same extent as they do with real conspecifics, provided these virtual objects mimic the characteristic kinetics of zebrafish swim bouts. Other naturalistic cues previously implicated in shoaling, such as fish-like shape, pigmentation pattern, or non-visual sensory modalities are not required. During growth, the animals’ stimulus preferences shift gradually, matching self-like kinetics, even in fish raised in isolation. Virtual group interactions and our multi-agent model implementation of this perceptual mechanism demonstrate sufficiency of kinetic cues to drive assortative shoaling, a phenomenon commonly observed in field studies. Coordinated behavior can emerge from autonomous interactions, such as collective odor avoidance in *Drosophila*, or from reciprocal interactions, such as the codified turn-taking in wren duet singing. We found that individual zebrafish shoal autonomously without evidence for a reciprocal choreography. Our results reveal individual-level, innate perceptual rules of engagement in mutual affiliation and provide experimental access to the neural mechanisms of social recognition. (239/250 words max)

**Significance Statement:** Social affiliation is ubiquitous in the animal kingdom, but fundamental sensory cues driving group formation remain elusive. During swarm behavior, for example, individuals dynamically exchange sensory cues with their neighbors, presenting an intertwined choreography opaque to formal analysis of causal stimulus-response relationships. Using a virtual interaction assay for psychophysical analysis, we solved this issue and identify biological motion as the irresistible trigger of social affiliation in zebrafish, *Danio rerio*. Given that many species form groups including shoals, flocks and herds, perceptual mechanisms of social recognition and their underlying neural circuits are likely shared across vertebrates. The identification of fundamental affiliation-inducing cues is a prerequisite for relating individual-level sensory-motor transformations to collective behavior. (112/120 words max)

## Main Text

Social interactions are essential to animals for survival and reproduction, suggesting that dedicated neuronal circuits exist to process socially relevant information (1–3). Indeed, specific groups of neurons in flies, mice and primates exert causal roles on social tasks such as social recognition, affiliation, mating and aggression (4–6). Neuromodulators such as oxytocin, serotonin and tachykinin regulate these behaviors across species and provide striking examples of evolutionary conservation in the control of social interactions, highlighting the potential of studying fundamental social principles in model organisms (4, 5, 7).

Social behaviors are triggered and regulated by conspecific cues and identifying their precise nature is central to social neuroscience. The best understood examples of such cues are pheromones which control innate behaviors via streamlined olfactory circuits (4, 8, 9). In *Drosophila melanogaster*, for example, the pheromone 11-cis-vaccenyl acetate activates olfactory sensory neurons which express the olfactory receptor Or67d. Output from Or67d neurons triggers sex-specific changes in mating behavior via sexually dimorphic connections with brain regions such as the lateral horn (9). In comparison, much less is known about fundamental visual cues, despite the fact that vision is required for many social behaviors including collective motion in groups (10), particularly in humans where the importance of pheromones is unclear (8). Studies on display behavior in cichlids and face processing in primates provide just a glimpse of socially relevant visual cues awaiting discovery (1, 11, 12). One major challenge to analyzing vision during social interactions is the dynamic, intermingled nature of visual cues exchanged by interacting animals. This hampers a systematic analysis of stimulus-response relationships and each animal′s causal contribution to the joint behavior, particularly in animal groups engaged in collective behavior such as swarm coordination (3, 13, 14). Quantitative descriptions of freely moving fish shoals, bird flocks and human crowds provide evidence that formation and maintenance of intraspecific groups are governed by a small number of simple behavioral rules such as long-distance attraction and short-distance repulsion among individuals (3, 10, 15–17). However, the challenge of isolating the fundamental visual cues and perceptual processes driving collective behavior remained unsolved and prevented dissecting the mechanistic implementation of collective rules at the level of neural circuits.

A powerful model of collective behavior is zebrafish shoaling, a form of affiliation with conspecifics that facilitates predator avoidance, foraging and stress coping (3, 18, 19). Mutual attraction among zebrafish develops between ten to twenty days of age when an increasing fraction of swim steering events become socially biased towards neighboring fish and animals maintain a preferred distance from one another (17, 20, 21). Previously, sensory triggers of shoaling were analyzed in adult fish by recording the location of a focal individual relative to one or several test fish separated by a transparent vertical divider. Such experiments revealed, that individual animals were more attracted towards larger groups than towards single animals and preferred pigmentation types experienced during early development (22, 23). More recently, analysis of adult zebrafish responding to semi-realistic fish images on computer screens or biomimetic replica revealed effects of stimulus size, color, shape and motion on attraction (14, 24, 25). Together, these results paint a complex picture of multiple interacting cues without any one visual feature dominating attraction. Bridging the gap between shoaling rules inferred from freely moving animals and attraction towards stimuli across a divider requires dynamic stimulus presentation and analysis of unrestrained interactions with artificial stimuli. Such experiments become feasible through the advent of closed-loop stimulus presentation and observer-centric virtual reality techniques (26, 27). In this study, we use a virtual-reality setup to present dynamic social stimuli to freely swimming animals at elevated throughput for psychophysical analysis in comparison to natural shoaling with real conspecifics. We find that kinetics features representing fish-like biological motion rather than photorealistic appearance trigger complete and persistent unfolding of shoaling. We analyze shoaling of developing juvenile animals with stimuli that cover a range of kinetic parameters and find an age-specific preference for self-like motion. By comparing mutual interactions between fish to interactions with non-interactive stimuli and simulated interactions in a multi-agent model, we propose that individuals shoal autonomously without reciprocal dialog. Together, these results outline perceptual principles of social recognition and provide a starting point to analyze the organization of neural circuits controlling collective behavior.

## Results

To investigate social affiliation, we tracked pairs of fish freely swimming in shallow dishes (Fig. 1A). Under such conditions, zebrafish readily engage in shoaling (20). Fig. 1B shows an example of two juvenile fish (26 days post-fertilization, dpf) spontaneously following each other closely over almost the entire observation period of 20 minutes. The inter animal distance (IAD, 5-20 mm) is well below the average distance expected from chance encounters (40 mm) (Fig. 1B), a hallmark of shoaling (20). We defined attraction as the percent reduction in mean IAD relative to control IAD expected by chance; this index was highly significant in each of 7 pairs at this age (Fig. 1B). In control experiments, we found that fish swimming individually in vertically stacked transparent dishes were still highly attracted to one another, indicating that vision is sufficient for shoaling (Fig. S1). Vision was also required: upon turning off all visible light, animals separated within seconds to chance level IAD (Fig. S1).

**Fig. 1:**
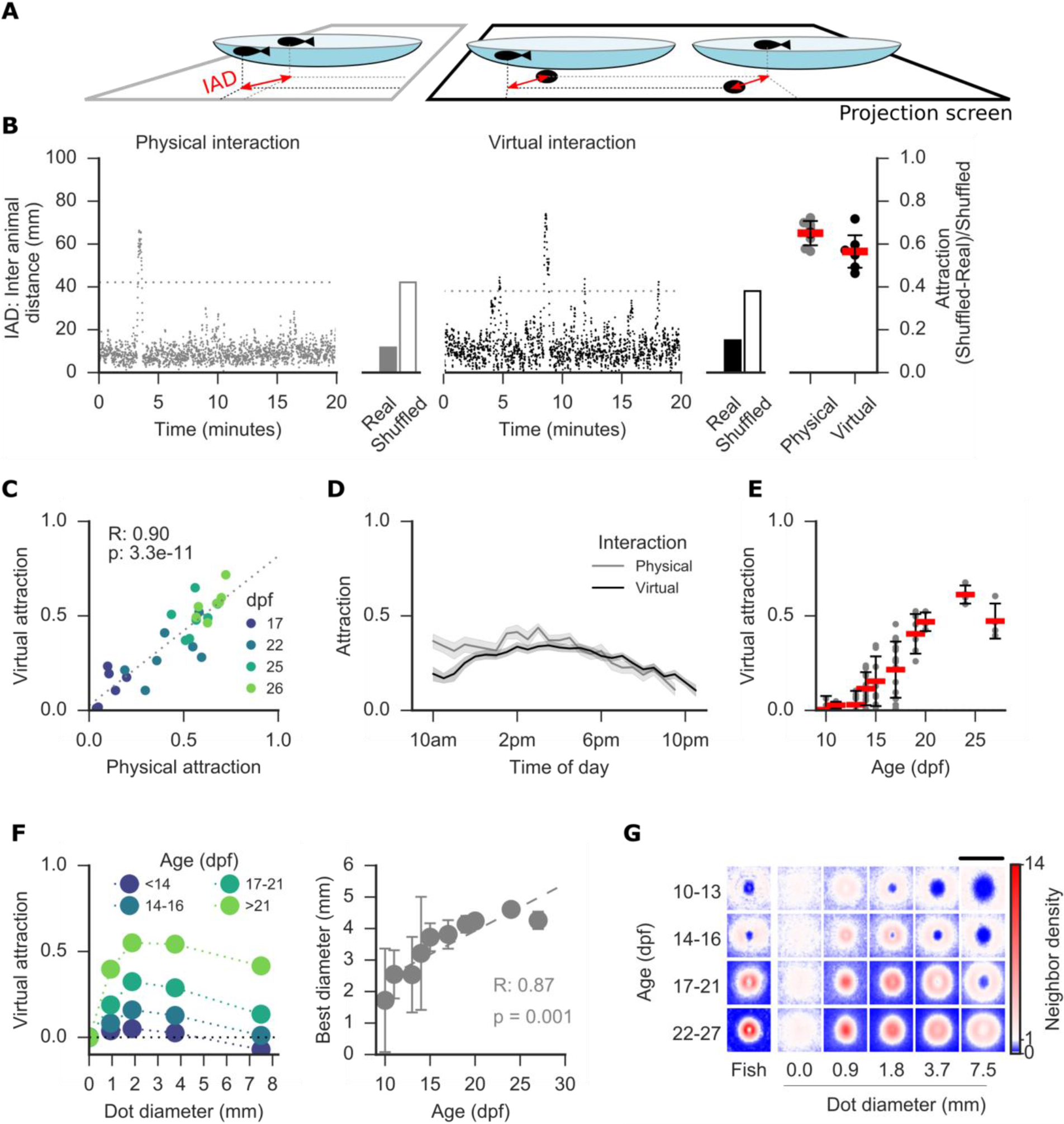
Virtual interactions driven by cross-projected interactive dots. **(A)** Schematic of experiments: Two animals interact physically (left) or virtually (right) in a watch glass of 10 cm diameter. In the virtual condition, a black dot is projected onto a screen below each dish at the location of the animal in the other dish. Double arrows indicate dot-animal distance or inter animal distance (IAD). **(B)** Representative IAD traces for one pair tested successively in both interaction modes. Bar graphs indicate mean real IAD_r_ during this period vs. chance level IAD_s_ obtained by time shuffling the position data. Right: An attraction index is calculated as (IAD_s_- IAD_r_)/IAD_s_. Age: 26 dpf. Data points indicate individual pairs, red bars indicate mean +/− 1 SD. **(C)** Attraction in individual pairs correlates between interaction modes across age. N=7 pairs per age group. **(D)** Attraction persists throughout the day. Traces represent mean, shading represents 95% CI. N=7 pairs (physical), 21 pairs (virtual), age: 23-25 dpf. **(E)** Attraction in the virtual mode matures between 2-3 weeks of age. Data points represent individual pairs. Red bars indicate mean at each age +/− 1 SD. N=70 pairs. **(F)** Most attractive dot diameter increases with age. Left, data points represent mean attraction for four age groups. Right, data points represent mean of best diameter over animals at each age +/− 1 SD. Dashed line represent linear fit through the mean values. Same data as (E). **(G)** Neighbor density distribution during physical interactions with another animal (left) and virtual interactions via black dots of variable diameter (right). A focal fish defines the center of each map. Red and blue color indicate higher and lower probability than chance, respectively, of finding the neighbor animal at a given location. Maps are mean probability over multiple animals. Physical interaction data (Fish) are the same as in panel C. Virtual interaction data are the same as in panel E. Scale bar represents 60 mm.

These findings prompted us to devise a simple virtual reality shoaling assay to reveal the fundamental visual cues that drive shoaling. We placed fish in separate dishes above a projection screen onto which we cross-projected in real time a black dot at the location of another fish (Fig. 1A). This dot virtually links two physically separated fish and was therefore termed ′interactive′. Notably, it enables mutual interactions in the absence of visual detail such as body shape, pigmentation, tail motion, depth, or texture, cues previously implicated in regulating shoaling (14, 24, 26). Pairs readily interacted via interactive dots of 3.7 mm diameter (Fig. 1B); their attraction was on average 87% of physical attraction measured in the same animals (Fig. 1B). The strength of attraction increased sharply with age and was correlated between physical and virtual conditions from 17 to 26 days of age, suggesting that both assays probe a related behavior that matures during this time (Fig. 1C). Shoaling is a persistent behavior of zebrafish in the wild(19). Accordingly, physical and virtual attraction were maintained over a 12 hour period and fell only towards the evening (Fig. 1D), demonstrating persistent engagement of the animals with interactive dots.

Onset of shoaling occurs at around 2 weeks of age (20, 21), a period of rapid development in zebrafish, including a doubling in body length from 4 mm to 8 mm within 2 weeks (28), and a gradual transition from well-isolated swim bouts to near-continuous swimming (28, 29) (Fig. S3). To dissociate the roles of increased social drive and animal size on mutual attraction during animal growth, we modulated the dot diameter from 0.9 mm to 7.5 mm and recorded virtual interactions in 70 pairs from 10 to 27 dpf (Fig. 1E-G). From 14 dpf, an increasing fraction of pairs showed attraction to 3.7 mm dots reaching 100 percent of pairs at 19 dpf (Fig. 1E). Attraction was strongest to dots of diameters between 1.8 and 3.7 mm and we detected a positive correlation between age and the most attractive dot diameter ranging from 3 mm at 14 dpf to 4.5 mm at 24 dpf (Fig. 1F). These sizes correspond to the parts of a juvenile zebrafish that provide the highest contrast such as the head and the eyes.

Natural shoaling is explained in part by two behavioral rules: Long-range attraction and short-range repulsion together result in characteristic animal spacing (3, 17). Neighborhood maps representing the likelihood of finding a neighbor in space reveal a time average of these opposing behaviors (16). We compared neighborhood maps of virtually versus physically interacting animals with respect to this signature. Physically interacting animals less than 14 dpf mainly exhibited repulsion and attraction was increasingly prominent in older animals (Fig. 1G). Maps of virtually interacting animals were most similar to physical interaction at dot sizes of 1.8 to 3.7 mm with a ring of attraction around a central zone of repulsion (Fig. 1G, S4). Short-range repulsion from the most attractive stimuli indicates that dot stimuli trigger shoaling behavior rather than pursuit of potential prey. From these results, we conclude that a circular black dot can act as a visual cue sufficient to drive shoaling provided it interactively mirrors the position of another animal.

Next, we sought to understand the minimal motion parameters that render a dot stimulus attractive to induce shoaling. Object speed is a key stimulus feature in hunting and escape behavior (30–32), suggesting dot speed might also regulate social affiliation. Alternatively, animals might extract higher order motion parameters such as acceleration or path curvature. For example, acceleration oscillates once every swim bout because propulsion and gliding alternate at about 1-2 Hz (29), a kinetic signature of biological motion (33) in zebrafish that may be used to detect conspecifics.

To address these possibilities, we measured individual animals′ attraction to dots moving along a set of synthetic stimulus paths that dissociated swim kinetics and swim paths by replacement with circular paths and constant speed, respectively (Fig. 2A). Such dots were non-interactive, moved identically for each animal and were, hence, called ′passive-attractive′. Fish also followed passive-attractive dots allowing us to quantify individual′s unilateral attraction towards such dots, in contrast to bilateral attraction measured in pairs mutually interacting via cross-projected dots (Fig. 2B). We noticed a preference for dot movement with natural swim kinetics over movement at constant speed that was largely independent of the dot path (Fig 2B), suggesting that fish recognize swim kinetics as a sign stimulus (11) for social affiliation.

**Fig. 2:**
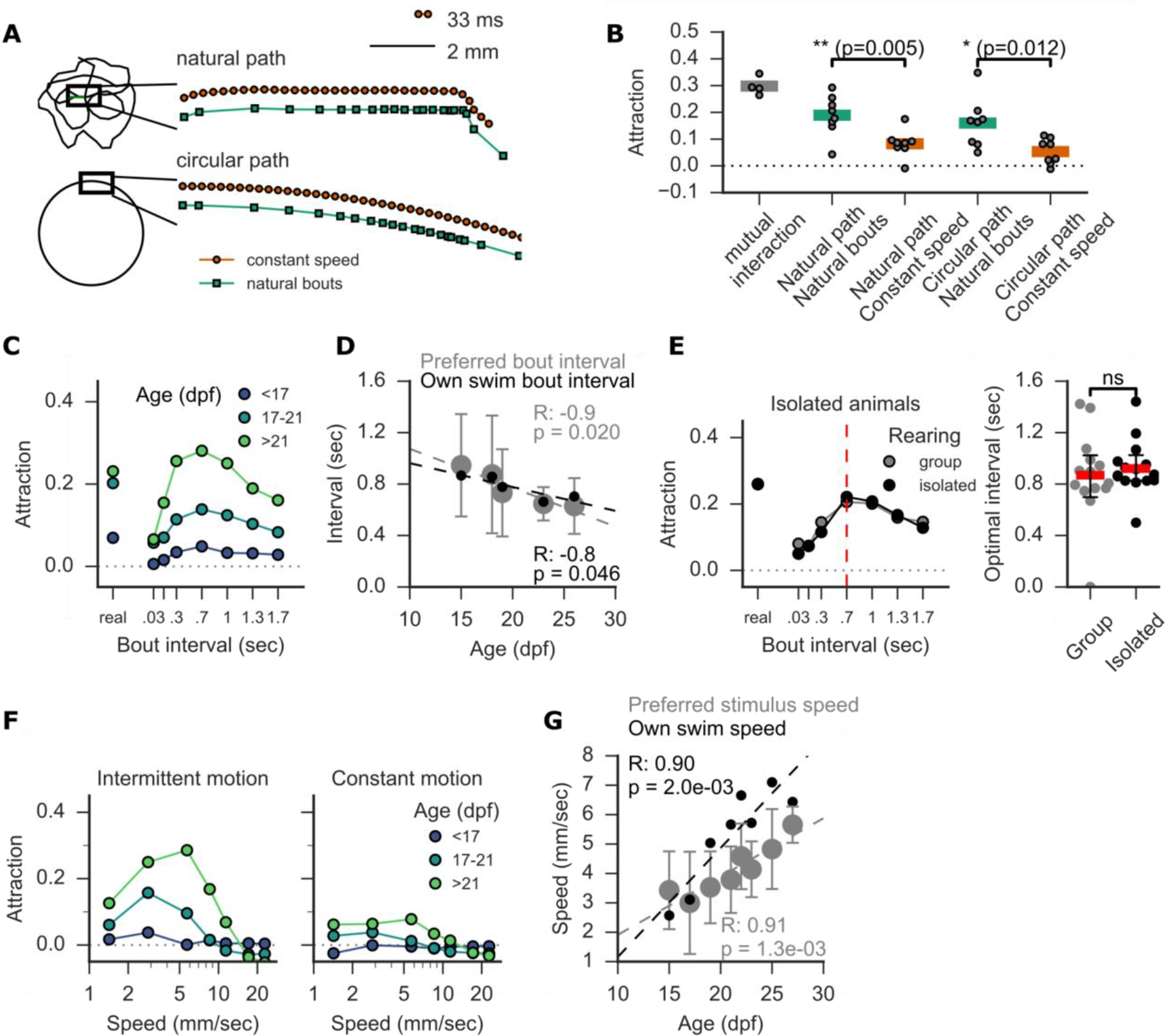
Fish-like stimulus kinetics trigger attraction towards passive-attractive dot stimuli. **(A)** Schematic of stimulus paths and kinetics: Colored data points indicate dot position in consecutive frames (33 ms). Constant speed paths were generated by interpolating natural paths while maintaining average speed. **(B)** Natural bout kinetics are necessary and sufficient to render dot stimuli attractive. N=8 animals. **(C)** Attraction to dots moving intermittently along a knot-shaped path. Bout interval was varied at fixed average speed. Data points represent mean attraction over animals at each age group. N=15-30 animals per age group. **(D)** Optimal bout interval (gray) tracks age-specific spontaneous swim bout frequency (black). Error bars are 1 SD. Dashed lines represent linear fit through group means. Same data as (C). **(E)** Rearing animals in isolation does not affect overall attraction or optimal interval at 18 dpf. N=15 animals per group. NS=not significant. **(F,G)** Constant speed stimuli yield low attraction at all speeds. Intermittent motion stimuli are most attractive at speeds that parallel natural swim speed across age 15-27 dpf. Gray data points represent mean optimal speed +/− 1 SD. Black data points represent average swim speed in the absence of a stimulus, n=15-30 animals per age group. Dashed lines represent best linear fit through group means.

Next, we decomposed natural swimming by simplifying its bout structure. We measured attraction towards dots moving in discontinuous jumps (bouts) over a range of intervals from continuous to intermittent bout-like motion at a fixed average speed. Attraction was strongly regulated by bout interval from 0.03 s to 1.3 s (Fig. 2C). We detected a negative correlation of the preferred bout interval with age: it ranged from 1 s to 0.6 s between 15 - 27 dpf, closely tracking the fish′s own spontaneous swim bout frequency at each age (Fig. 2D). Strikingly, the overall attraction and age-specific optimal bout interval of fish reared in isolation was indistinguishable from control animals raised in groups (Fig. 2E). This indicates that an innate mechanism underlies the development of stimulus pattern recognition.

To ask how speed modulates attraction, we presented fish with intermittent and continuous motion at average speeds of 1.5 to 24 mm/s. Intermittent motion was more attractive at all speeds (Fig. 2F), and attraction dropped sharply at speeds above 10 mm/s. The preferred dot speed rose from 3 to 6 mm/s between 15 and 27 dpf, again tracking the animal′s own spontaneous swim speed (Fig. 2G). In contrast, continuous motion was ineffective at all speeds, as was inverting dot contrast to light on dark (Fig. S5). This tuning to self-like motion is further evidence that virtual interactions represent social affiliation.

Juvenile zebrafish perform swim maneuvers on a sub-second time scale resulting in rapidly fluctuating IAD time series of individual pairs (see Fig. 1B). To ask on what time scales fish integrate visual information to assess dot attractiveness, we analyzed changes in IAD around transitions of passive-attractive dot stimuli between intermittent and continuous motion (Fig. 3A): On average, fish had low steady-state IAD during intermittent motion and high IAD during continuous motion (Fig. 3B). After a transition from intermittent motion to continuous motion, IAD rose within 2 seconds and approached the higher steady-state within one minute (Fig. 3B). The increase in IAD reflects a combination of two processes: (i) the animal re-evaluating the stimulus, potentially with memory of stimulus history; and (ii), transition to a new steady-state IAD at a rate depending on swim speed and arena geometry.

**Fig. 3:**
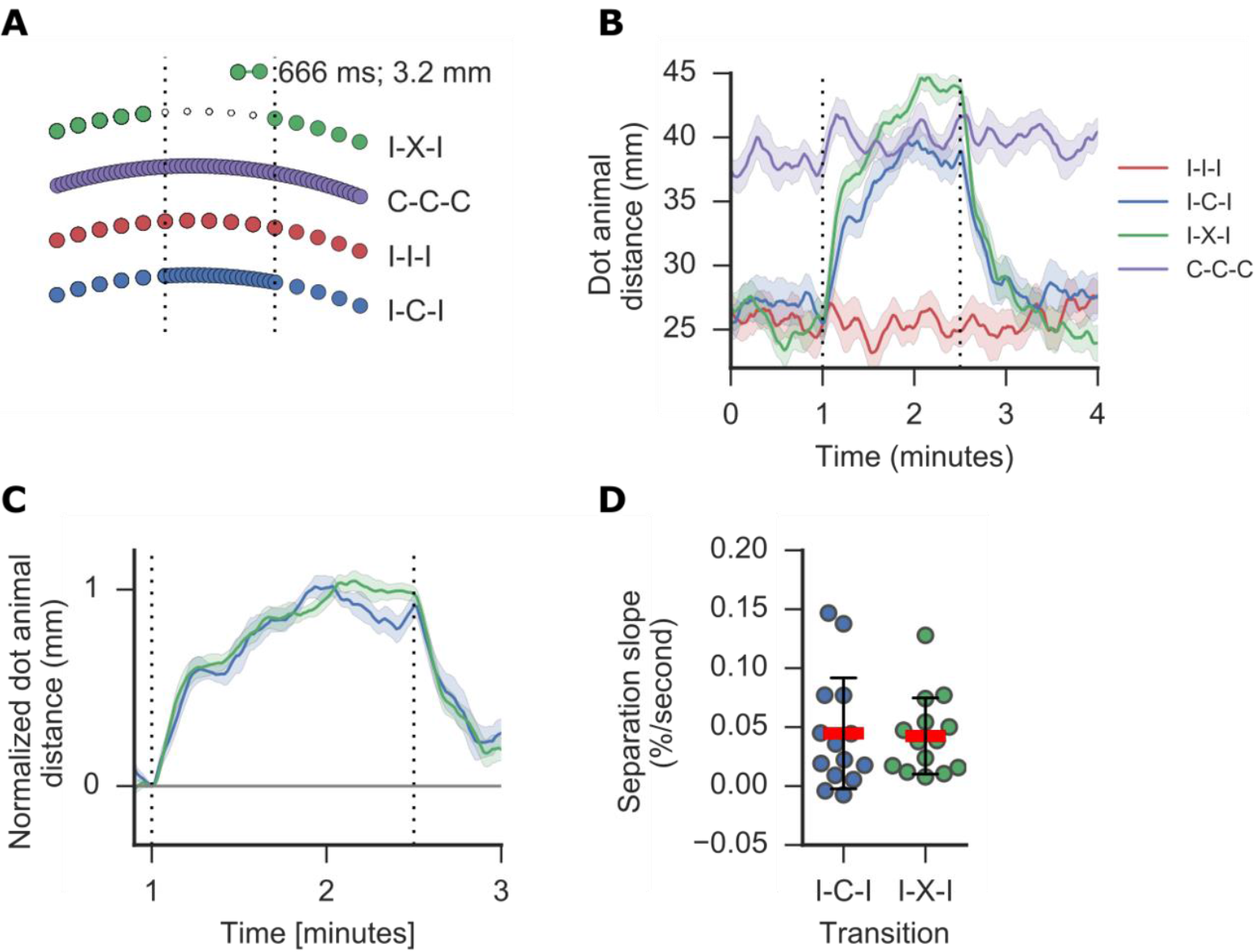
Social attractiveness is evaluated instantaneously. **(A)** Dot attractiveness is modulated at transitions from intermittent motion (I) to constant speed (C) or an invisible dot (X). **(B)** Dot-animal distance reaches a steady state within one minute after the transition (dashed lines). This separation represents a combination of a change in attraction and drift towards a new steady state. Lines represent mean distance, shading represents 95% CI, N=14 animals, 20-23 dpf. **(C)** Same data as B, normalization highlights similarity of the time courses. **(D)** Slope of dot-animal distance during 5 seconds after stimulus transition.

To isolate the rate of stimulus re-evaluation, we compared separation from a dot transitioned from intermittent motion to continuous motion versus separation from a dot turned invisible. Necessarily, animals fully devalue an invisible dot instantaneously. Animals separated at similar rates from invisible or continuously moving dots (Fig. 3B) and normalizing IAD time series of separation to account for different steady-state levels rendered the rates indistinguishable (Fig. 3C, D). This implies that juvenile zebrafish instantaneously evaluate kinetic attractiveness without memory of stimulus history.

A fundamental challenge in analyzing mutual interactions is dissecting each individual′s contribution. For example, mutual attraction within a pair and individual′s apparent sociability may be determined by each individual′s social drive towards the partner, by each individual′s attractiveness as evaluated by the partner, or a combination. In addition, mutual interactions may require explicit reciprocity with the partner, for example in the form of codified turn-taking as observed in duet singing in wrens. During zebrafish shoaling synchronization of swim bouts between neighbors (21) may provide a cue to boost attraction in a synergistic manner when such reciprocity is detected. To ask if individual′s social drive increases with reciprocating partners we sequentially measured bilateral attraction within all possible 105 virtual pairings of 15 animals via interactive dots. In each animal, we also measured unilateral attraction towards a passive-attractive dot (Fig 4A). Unilateral attraction is not confounded by the individual′s attractiveness, but instead is a direct reflection of social drive. Unilateral attraction of individuals and bilateral attraction within specific pairs were repeatable across trials but varied substantially between animals (Fig. 4B and S6). Mutual attraction of individuals to other animals was correlated across pairings, resulting in a range of mean bilateral attraction (MBA) reflecting apparent sociability of individual animals from effectively non-social to highly social (Fig. 4b).

**Fig. 4:**
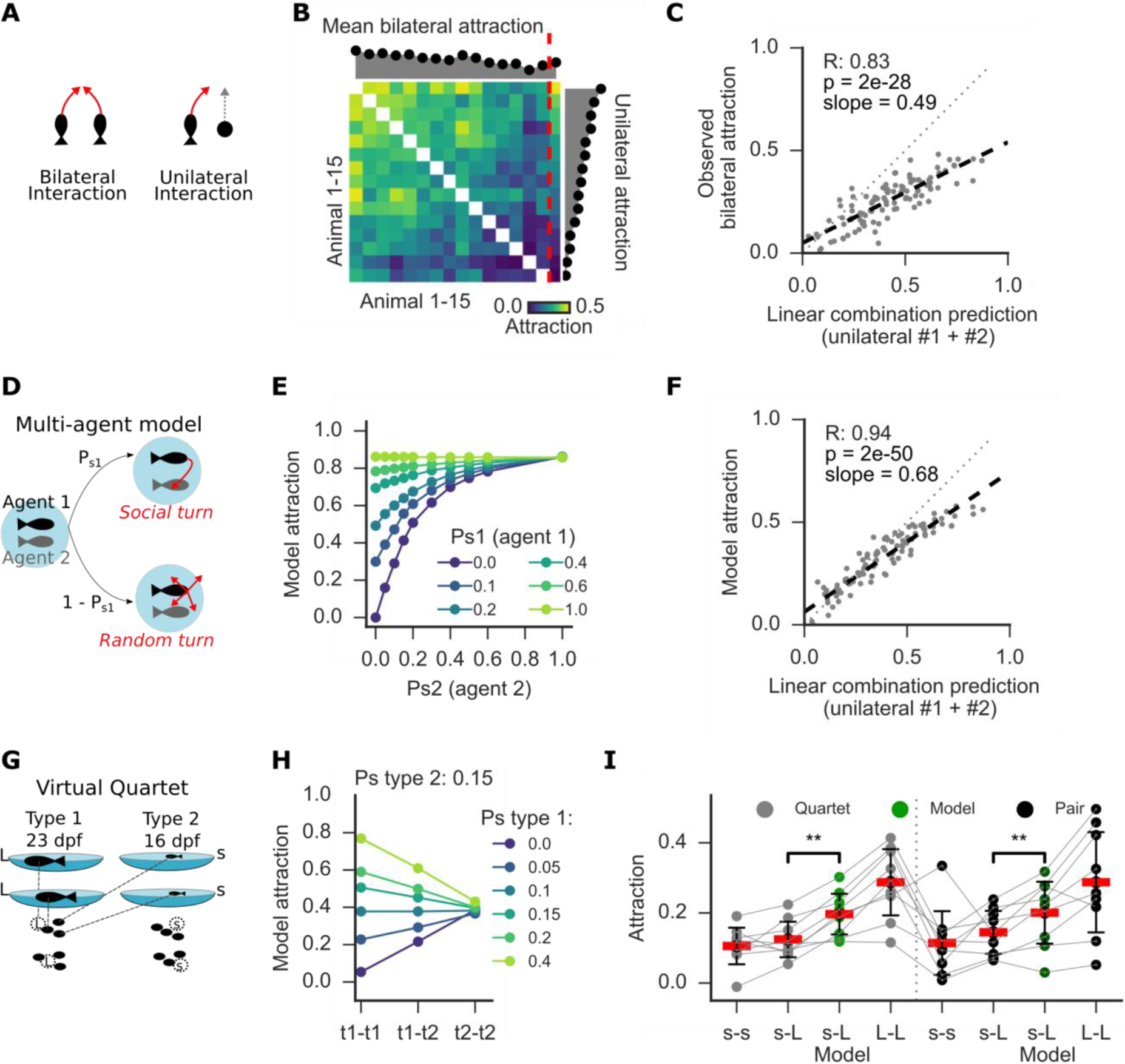
Individual social drive and age-specific kinetic preference predict mutual interactions. **(A)** Both animals potentially contribute social drive and attractiveness to mutual attraction observed in bilateral interaction. A single animal′s social drive determines observed attraction during unilateral interactions. **(B)** 15 animals are sequentially linked for bilateral interactions via interactive dots in all possible 105 pair-wise combinations. Each animal is also exposed to a passive-attractive dot moving intermittently to measure unilateral interaction. Rightmost column separated by red dashed line and histogram right represent unilateral attraction and determines plot order. Colors represent mean attraction for each pair over six repeated trials. Histogram above indicates column-wise mean attraction. **(C)** Pair-wise bilateral attraction is predicted as the sum of the two individuals′ unilateral attraction. Each data point represents one of 105 possible pairings. Dashed line is linear fit through data. Dotted line is unity. **(D)** Schematic of multi-agent model inspired by Hinze et al.(20). Agent n turns towards other agents with social turn probability P_sn_ or into a random direction with probability 1-P_sn_. Agent speed, bout rate, arena size and simulated frame rate match our data. **(E)** Mutual attraction between agents varies with social turn parameter P_sn_. P_s1_ = 0 corresponds to unilateral attraction of agent 2. See methods for details on this model. **(F)** Analogous to panel C, bilateral attraction observed in the model was predicted by a linear sum of unilateral attraction. Each dot represents one of 105 models using p_sn_ parameters corresponding to the animal pairs in panel C. Dashed line represents unity. **(G)** Schematic of a virtual quartet consisting of two animals each of two types (16 dpf and 23 dpf). Each animal sees black dots underneath at the location of the other three animals. All dots are of the same size. **(H)** Multi-agent model for quartet of 2 types differing only in P_sn_ predicts attraction between types (t1-t2) as the mean of attraction within types. **(I)** Attraction between types(s-L) was lower than predicted by the model (green). This preference was similar when tested simultaneously in a quartet or separately in pairs. Data points represent mean attraction over 4-6 repeated trials. s-L attraction represents mean over the four possible s-L combinations. Red bars indicate group means +/− 1SD. N=9 × 4 animals. Gray lines connect repeated measures of the same pair. **: p<0.01, related sample t-test.

We found that unilateral attraction correlated with MBA (R=0.91, Fig. 4B) implying that mainly social drive, and not attractiveness, determines mutual interactions and, thus, apparent sociability in our data. We could therefore predict bilateral attraction for each pair as the linear sum of each individual′s unilateral attraction (Fig. 4C).

A multi-agent model implementing a simple attraction rule (20) controlled by a social drive parameter recapitulated this linear prediction in the experimentally observed range of attraction values (Fig. 4D-F). Importantly, this model does not explicitly implement synergies in mutually interacting pairs. Thus, juvenile zebrafish did not distinguish between interactive versus passive-attractive dots. We conclude that individuals autonomously evaluate conspecific motion as a shoaling stimulus, solely as a function of their own social drive.

Field observations of fish shoals commonly describe affiliation preferences for conspecifics and size-matched animals (3). Such assortative shoaling can help individuals to evade detection by confusing predators who may target rare phenotypes, a form of selective predation also known as the oddity effect (3). To ask how differences in social drive and preferences for specific motion kinetics might act to sort larger groups, we analyzed virtually interacting quartets composed of two younger, weakly social fish (16 dpf) and two older, strongly social fish (23 dpf)(Fig. 4G). Each animal saw three dots of equal size at the position of the other fish. This configuration discards stimulus size as a factor influencing affiliation, but retains age-specific stimulus kinetics. To test for affiliation preferences within a quartet, we calculated pairwise attraction within and between the age groups. The multi-agent model predicts precisely intermediate attraction between groups compared to attraction within each group in the absence of any age-related preferences (Fig. 4H). As predicted, we found weakest and strongest attraction within the young and old groups, respectively (Fig. 4I). However, attraction between age groups was lower than predicted by the simple model as the mean of the two age groups (Fig. 4I). This result is consistent with active selection of preferred stimulus kinetics among multiple stimuli. To ask if indirect effects of different swim kinetics can explain assortative shoaling (34), we extended the model to implement group differences in swim speed, bout interval and an explicit within-group preference as a proxy for kinetic preferences. Of these parameters, only explicit within-group preference reproduced the observed affiliation preference (Fig. S7). For comparison, we virtually linked the six possible pairs of each quartet for pair-wise interaction. Attraction within and across age groups was indistinguishable between pairs versus quartet, demonstrating that the presence of two additional stimuli neither confuses, nor enhances the animal′s preferences for motion kinetics (Fig. 4I). From these results, we conclude that age-specific motion cues are sufficient to drive assortative shoaling.

## Discussion

Until now, sensory cues implicated in shoaling were numerous and their link to naturally unfolding behavior was unclear. Our results suggest that the perceptual basis of shoaling is simpler than previously thought. We discovered that the social ′instinct′ (11) of zebrafish is released instantaneously by visual features of the swim kinetics of another zebrafish. Imposing the bout structure typical of a juvenile fish on the movement of a projected, two-dimensional dot was sufficient to elicit the full suite of shoaling behaviors in a virtual reality arena.

Biological motion is a potent releaser of intraspecific and interspecific interactions in fish (25, 35), birds (36) and mammals including humans (37), and impaired interpretation of biological motion has emerged as an early heritable marker for autism (38, 39). In human psychophysics, biological motion typically refers to motion of body parts relative to each other, which is readily recognized by humans, even when the position of each limb is indicated only as a dot on a light point display (33). While naturalistically moving dots arrangements still represent a complex stimulus, local detection of acceleration that is consistent with biological agents emerges as one fundamental ′life-detector′ evaluating such stimuli (37). Our results suggest that a related detector functions as a core perceptual mechanism whose activation readily drives persistent shoaling in zebrafish.

Social cues often elicit innate behaviors, and social experience can profoundly shape such innate responses, providing insights into mechanisms of development and plasticity. One powerful manipulation, for example, is social isolation. In mice, solitary males, but not socially housed males, launch attacks on intruder animals (40). In contrast to this modulation, we found that preference for age-specific self-like motion formed indistinguishably in socially reared and isolated juveniles suggesting a fully innate developmental mechanism. This raises the possibility that juvenile zebrafish match visual cues against a cognitive representation of idiosyncratic, self-like biological motion which might be generated from proprioceptive feedback or an efference copy of swimming. Alternatively, the visual system might generate a gradually developing motion template independent of self-motion.

Coordinated behavior can emerge from autonomous interactions, such as collective odor avoidance in *Drosophila*, where mechanosensory interactions upon animal collisions enhance the response probability to escape from mildly noxious stimuli (41). Social interactions can also require explicit reciprocity, such as the codified turn-taking in wren duet singing, which is reflected in neural encoding of the jointly produced song by each individual (42). We demonstrate persistent attraction of individual juvenile zebrafish towards non-interactive stimuli. Naturally, the observed affiliation of two agents who both move towards each other is closer than affiliation observed between one individual and a non-interactive agent. Within-animal comparisons of bilateral and unilateral interactions together with our multi-agent model suggest that individuals equally shoal with passive-attractive and interactive stimuli. We conclude that juvenile zebrafish autonomously evaluate and respond to motion cues during shoaling rather than coordinating an explicitly reciprocal behavior. This lack of reciprocity suggests that simple passive-attractive stimuli may suffice to activate neural circuits for social processing in restrained animals during functional imaging.

Our analysis revealed consistent variability in mutual attraction between animal pairs which we assigned largely to differences in individuals′ responsiveness to motion cues, the social drive of each animal. Recent studies specifically focused on inter-animal differences in repeated measurements of behavior to reveal genetic and neural causes of phenotypic variability (39, 43, 44) and its effect on collective behavior (34). It will be important to analyze individual animals over longer periods of time to determine persistence and heritability of different social personality types. We speculate that individual differences in the perception of motion cues partially predict social drive, consistent with impaired interpretation of biological motion in children with autism (38). In addition, internal states set by neuromodulatory systems shape behavioral individuality (43, 44). Differences across these domains are, in principle, detectable at the level of neural activity and provide exciting opportunities to study mechanistic causes and social consequences of phenotypic diversity.

Defining minimal visual social cues in a genetic model organism opens the door for future studies on the underlying brain mechanisms, analogous to the discovery of pheromones and their role in shaping animal behavior (4, 8, 9). Our results provide a baseline set of stimuli at the earliest social stage to which juvenile zebrafish likely add as they mature. Future work in older animals will reveal developmental stages when other naturalistic cues such as the conspicuous zebrafish pigmentation gain influence on shoaling decisions. In the meantime, our stimulus provides a clear path forwards for identifying those areas in the zebrafish brain tuned to biological motion stimuli displayed to restrained animals during functional imaging of whole brain activity and optogenetic interrogation (45–47). It will be interesting to trace the neural pathways that underpin the social instinct, from the detection of conspecifics by the visual system to the innate responses encoded in the hypothalamic and limbic centers of the forebrain. Since social affiliation is a common feature across the vertebrate taxon, the core principles of neural architecture are likely to be conserved from fish to human.

## Acknowledgments

We thank Marco Dal Maschio and Joseph Donovan for help with designs for the setup and statistics. We thank X. Jin, S.W. Flavell, D. Ventimiglia, S.J. Rahi, A. Tallafuss, P. Washbourne and members of the Baier laboratory for comments on the manuscript and discussions.

## Author contributions

J.L. and H.B. conceived the study. J.L. designed and conducted experiments and analyzed the data. J.L. and H.B. interpreted the data and wrote the paper.

## Data and materials availability

Analysis code is accessible at GitHub (link active upon publication). Setup design, control software and raw data will be made available upon request.

## Supplementary Materials

Materials and Methods

Supplementary Text

Figs. S1 to S7

Captions for Movies S1 to S2

